# Deep Learning Based Models for Preimplantation Mouse and Human Development

**DOI:** 10.1101/2024.02.16.580649

**Authors:** Martin Proks, Nazmus Salehin, Joshua M. Brickman

## Abstract

The rapid growth of single-cell transcriptomic technology has produced an increasing number of datasets for both embryonic development and *in vitro* pluripotent stem cell derived models. This avalanche of data about pluripotency and the process of lineage specification has meant it has become increasingly difficult to define specific cell types or states and compare these to *in vitro* differentiation. Here we utilize a set of deep learning (DL) tools to integrate and classify multiple datasets. This allows for the definition of both mouse and human embryo cell types, lineages and states, thereby maximising the information one can garner from these precious experimental resources. Our approaches are built on recent initiatives for large scale human organ atlases, but here we focus on the difficult to obtain and process material that spans early mouse, and in particular, human development. Using publicly available data for these stages, we test different deep learning approaches and develop both a model to classify cell types in an unbiased fashion and define the set of genes required to identify lineages, cell types and states. We have used our predictions to probe pluripotent stem cell models for both mouse and human development, showcasing the importance of this resource as a dynamic reference for early embryogenesis.

## Introduction

Mammalian development begins at fertilization, producing a single totipotent zygote from which the entire embryo and supporting structures emerges. The zygote then undergoes a series of reductive cleavage divisions, during which the embryo increases in cell number while maintaining its overall size. Although maternal RNAs are deposited into the oocyte, they have little influence on differentiation as zygotic transcription begins at relatively early stages, with zygotic genome activation (ZGA) occurring in mouse at the two-cell (2C) stage, and in human embryo at the eight-cell (8C) stage (Yuan et al., 2023). When the embryo reaches the morula stage (16C), the first lineage segregation occurs, where precursors of the placenta polarize and form the outer trophectoderm (TE) (Nishioka et al., 2009; Gerri et al., 2020). These surround the now internalized inner cell mass (ICM). As or shortly after (depending on the species) these two cell types are established, the embryo transitions to the blastocyst stage and produces a fluid-filled blastocoel cavity. During blastocyst maturation, a second lineage specification event is observed in ICM cells, as they differentiate to either Epiblast (EPI) or primitive endoderm (PrE). This is followed by the segregation of these lineages, such that the PrE becomes positioned between the EPI and blastocyst cavity. At this stage, the embryo is ready to hatch from the zona pellucida and implant into the uterine wall. During post-implantation development the TE gives rise to the placenta, PrE to the visceral and parietal endoderm, and EPI to the embryo proper (Gilbert, 2000; Saiz and Plusa, 2013; Riveiro and Brickman, 2020).

Since the explosion in single-cell sequencing techniques (scRNA-seq), this technology has been extensively applied to these accessible stages of embryonic development. Numerous datasets produced by different technologies have been used to attempt to understand these early cell fate choices (Biase et al., 2014; Borensztein et al., 2017; Boroviak et al., 2015; Chen et al., 2016; Deng et al., 2014; Fan et al., 2015; Goolam et al., 2016; Mohammed et al., 2017; Nowotschin et al., 2019; Posfai et al., 2017; Xue et al., 2013; Stirparo et al., 2021; Yanagida et al., 2022; Meistermann et al., 2021; Petropoulos et al., 2016; Xiang et al., 2019; Yan et al., 2013; Yanagida et al., 2021). Moreover, as stem cell technologies have been adapted to generate *in vitro* models for different aspects of early human development, these transcriptional profiles have been an essential resource for defining the developmental stage of *in vitro* cell types including human primed (Thomson et al., 1998) and naive embryonic stem cells (ESCs) (Gafni et al., 2013; Takashima et al., 2015; Bredenkamp et al., 2019a), TE stem cells (Dong et al., 2020; Cinkornpumin et al., 2020; Okae et al., 2018), PrE stem cells (Linneberg-Agerholm et al., 2019; Okubo et al., 2023) and various three-dimensional *in vitro* models of development (Kagawa et al., 2021; Liu et al., 2021; Yu et al., 2021; Yanagida et al., 2021; Fan et al., 2021). However, the need for manual isolation and the limited amount of high quality material compared to adult organ studies, has meant that obtaining large number of cells is difficult. In addition, the ethical challenges associated with obtaining human embryos means that information gained from even a limited number of cells is extremely valuable. As a result, there is a need to find coherent approaches to combine existing datasets and generate a useful and evolving tool that can be used to benchmark the ever increasing number of cell culture models and cell types. One approach to overcome these challenges and strengthen downstream analyses is to collate multiple scRNA-seq experiments together.

Traditional data integration techniques assume a linear relationship between datasets, which outperform non-linear approaches when batch effects are small and biological complexity low (Luecken et al., 2022). However, in early embryogenesis the myriad of diverse signaling, epigenetic and transcriptional events required to underpin regulative development makes it a robust but variable process. Moreover, individual sequencing techniques provide varying sequencing depths and different levels of technical noise which could drive the skewed integration of datasets. Lastly, computational demand scales linearly with the amount of data suggesting current approaches will soon become intractable (Angerer et al., 2017). These shortcomings have been addressed by deep-learning integration techniques which utilize neural networks and graphical processing units (GPUs) to collapse cells into lower dimensional latent space which is further used for downstream analyses (Lopez et al., 2018; Lotfollahi et al., 2021; Erfanian et al., 2023). These approaches have been already successfully applied in building numerous atlases such as *The Human Cell Atlas* (Regev et al., 2017), *The Tabula Sapiens* (Consortium* et al., 2022), *Human Lung Cell Atlas* (Sikkema et al., 2023) and several more recent organ or disease specific atlases (Swamy et al., 2021; Eraslan et al., 2022; Domínguez Conde et al., 2022; Suo et al., 2022). Despite promising results from these integrations, the underlying neural networks still suffer from a lack of interpretability. Although there have been some early attempts to do this, most have been difficult to interpret. For example, linearly decoded variational autoencoder (LDVAE) (Svensson et al., 2020), decouples the learned latent space into modules of linearly dependent genes, but like traditional data integration, suffers from the same linear assumption that similar cell states are coupled. Thus, while traditional cell type assignment is generally done by hand, these models could provide an excellent unbiased tool to assign cellular identity, but there are currently only limited approaches to distill the logic used by the underlying models.

In this paper, we employed state-of-the-art computational tools to build transcriptomic models of mouse and human preimplantation development. We collated single-cell transcriptomic datasets of mouse and human preimplantation embryos and, by taking advantage of scvi-tools (Gayoso et al., 2022) for probabilistic modeling of single-cell omics data, built cell type and time-point classifiers. Secondly, we overcome a critical disadvantage of ‘black box’ deep learning models by implementing a SHapley Additive exPlanations (SHAP) (Lundberg and Lee, 2017) algorithm to interpret the logic behind lineage classification. Finally, we display the utility of these models to classify lineages generated from *in vitro* differentiation of mouse and human stem cells. We anticipate the models will serve as a valuable resource for the community, providing an evolving model that can be used to probe phenotype and benchmark an increasing number of *in vitro* cell culture models.

## Results

### A comprehensive model for annotation and integration of preimplantation mouse and human embryos

To build a reference model we collected *in vivo* preimplantation scRNA-seq datasets for both mouse and human embryos covering the stages depicted in Fig. 1A. The datasets were selected only if they were part of a published peer-reviewed article and contained cell metadata for the time of collection and initial cell type annotations. These datasets were then filtered to select only wild-type embryos, rather than experimental genotypes or *in vitro* stem cells reported alongside these datasets. Based on these criteria we built a ground truth reference model. Thirteen mouse and six human datasets satisfied these criteria and represent eleven years of studies employing five different sequencing techniques (Table 1, S1A, S2B). The datasets containing the largest numbers of cells were Nowotschin et al. (2019) and Petropoulos et al. (2016), for mouse and human respectively (Fig. 1B, S1B, S2B).

**Figure 1.**
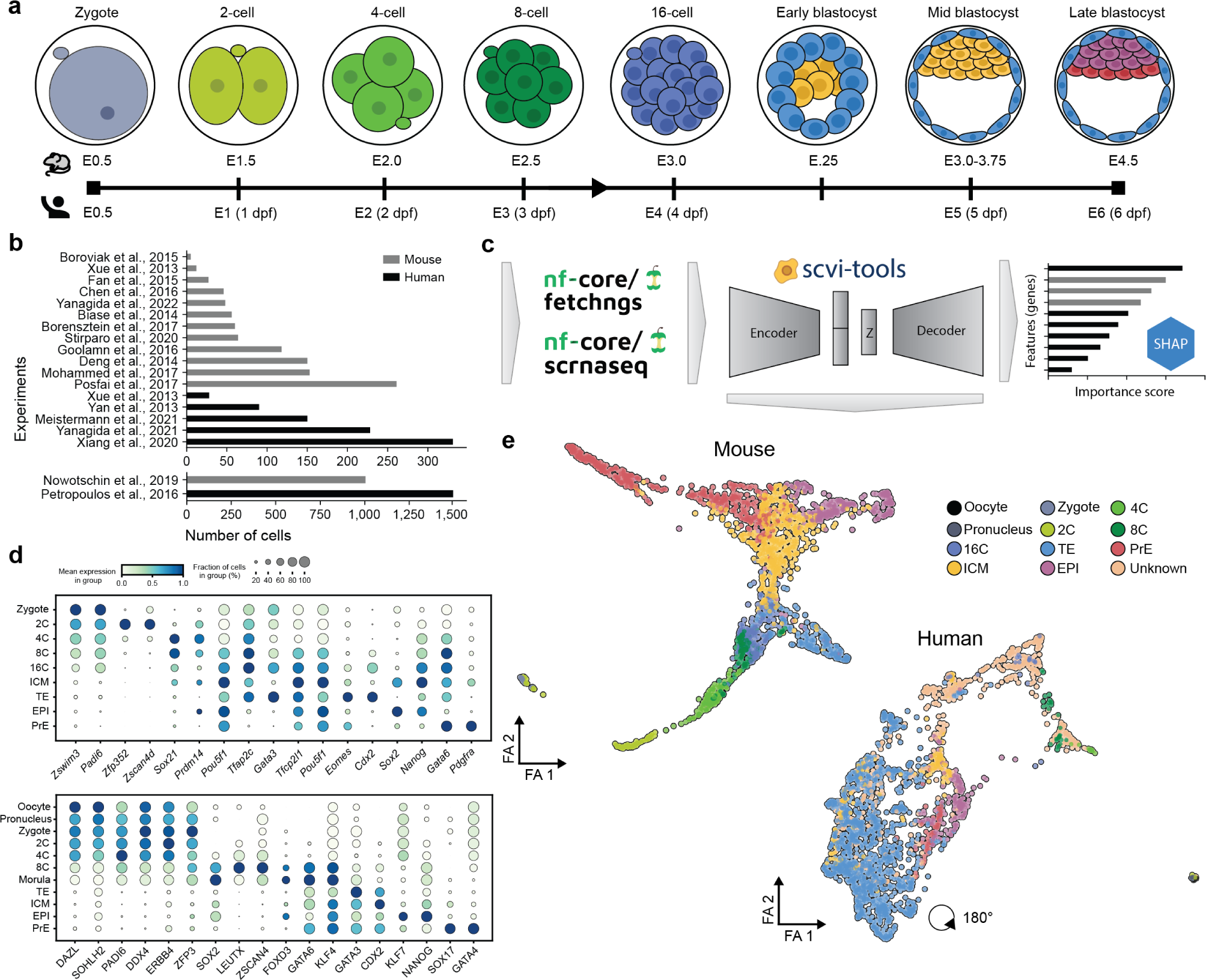
Summary of datasets used to build reference models. **a**) Schematic overview of mouse and human preimplantation development. **b**) Quantification of cells per publication which were collected for building the mouse (grey) and human (black) reference. **c**) Computational schematic of tools used to build and interpret the reference models. **d**) Gene expression of canonical markers for each developmental stage in mouse (top) and human (bottom) preimplantation. **e**) Reduced dimensional representation of preimplantation mouse (left) and human (right) datasets. dpf: days post fertilization, E: embryonic day

**Table 1.** List of published datasets used to train reference models.

| Organism | Publication | Technology | GEO Accession number |
| --- | --- | --- | --- |
| Mus Musculus | Biase et al. (2014) | SMART-seq | GSE57249 |
|  | Borensztein et al. (2017) | qRT-PCR | GSE80810 |
|  | Boroviak et al. (2015) | SMART-seq2 | E-MTAB-2958/2959 |
|  | Chen et al. (2016) | SMART-seq2 | GSE74155 |
|  | Deng et al. (2014) | SMART-seq 1/2 | GSE45719 |
|  | Fan et al. (2015) | SUPeR-seq | GSE53386 |
|  | Goolam et al. (2016) | SMART-seq2 | E-MTAB-3321 |
|  | Mohammed et al. (2017) | SMART-seq2 | GSE100597 |
|  | Nowotschin et al. (2019) | 10X v2 | GSE123046 |
|  | Posfai et al. (2017) | SMART-seq | GSE84892 |
|  | Xue et al. (2013) | qRT-PCR | GSE44183 |
|  | Stirparo et al. (2021) | SMART-seq2 | GSE159030 |
|  | Yanagida et al. (2022) | SMART-seq2 | GSE148462 |
| Homo sapiens | Meistermann et al. (2021) | SMART-seq2 | PRJEB30442 |
|  | Petropoulos et al. (2016) | SMART-seq2 | E-MTAB-3929 |
|  | Xiang et al. (2019) | SMART-seq2 | GSE136447 |
|  | Yan et al. (2013) | SMART-seq | GSE36552 |
|  | Yanagida et al. (2021) | SMART-seq2 | GSE171820 |
|  | Xue et al. (2013) | qRT-PCR | GSE44183 |

We placed an emphasis on automating the preprocessing by taking advantage of nf-core pipelines (Ewels et al., 2020) to download, align and quantify the datasets (Fig 1C). The entire process is version controlled and can be easily rerun to take advantage of improvements in aligner accuracy and performance, updated genome assemblies and gene annotations. Additionally, this setup allows for simple expansion with new *in vivo* datasets, and preprocessing can be run with the same exact parameters on *in vitro* datasets (query) for downstream benchmarking. To ensure our reference datasets are maintained, future iterations and their refinements will be versioned and accessible on Hugging Face (https://huggingface.co/).

Steps taken after preprocessing to gene transcript quantification diverged for the mouse and human datasets. Below we discuss the details of each of model and how we integrated the datasets. For each experiment, cell types were re-annotated in accordance to their developmental stages, while maintaining the original published annotations within the metadata. For example, *Deng et al.* (Deng et al., 2014) annotated early, mid and late stages of 2C cells, which we collapsed to 2C stage. Additionally, we discarded data arising from oocytes due to their low representation (N =13). Lastly, we identified the top 3,000 most highly variable genes (HVGs) between experiments for downstream analysis as subsetting analysis to HVG provides performance boosts for comparable cell type identification to that generated with the complete transcriptome (Wagner et al., 2012).

#### Mus Musculus

As the mouse dataset contained a mixture of full-length and UMI based single-cell sequencing, we normalized datasets generated using the SMART-seq1/2 protocol by gene length (Wagner et al., 2012; Phipson et al., 2017; Heumos et al., 2023). We discarded ribosomal, mitochondrial genes due to their possible contribution to variance and identification as highly variable genes, as well as *Ct010467.1* due to its high fraction of counts. Based on the quality control (QC) cells containing minimum of 20,000 transcripts per cell were retained (see methods). These steps yielded a final mouse dataset of 2,004 cells and 34,346 genes.

#### Homo Sapiens

In human, a number of datasets contained cell labels that were ambiguous or intermediate. To avoid introducing uncertainty for classifiers we set these labels to ‘Unknown’ despite the availability of annotation on time point and speculating possible identity. These cells were later used as an internal validation set during model optimization and to test our classifiers. Compared to the mouse dataset, the human dataset contains a disproportionate number of cells annotated as TE and although this can create an imbalance for our classifiers, we decided to retain all the cells. Given the scarcity of material we sought to bolster cell numbers for classification. Here, we reasoned a minimum of 15 cells per label would be required and so we collapsed all stages prior to the 8-cell stage to ‘Prelineage’. A similar collation was performed for all cells annotated as PrE regardless of developmental stage. Unlike the mouse studies, those of human preimplantation relied exclusively on full-read sequencing technologies. However, to ensure the model could be used to integrate cells sequenced using UMI-based technologies, we similarly transformed read counts by gene length.

#### Integration

Based on a previous study benchmarking integration strategies for scRNA-seq datasets (Luecken et al., 2022), we used single-cell variational inference (scVI) (Lopez et al., 2018) and scGen (Lotfollahi et al., 2019) to integrate existing datasets. We fined tuned parameters during training (Suppl. Table 1) using the *autotune* feature in scvi-tools. However, we observed that the overall best performance was achieved using 2 hidden layers and fitting to negative binomial distribution with early stoppage during training for both species. To assess the performance we tracked the evidence lower bound (ELBO) per epoch and calculated batch and biological conservation (S1C. S2C) using the scib-metrics package (Luecken et al., 2022; YosefLab/scib-metrics, 2023). scGen ranked first, but for mouse only. However, the tool was designed to integrate control and perturbed experiments which was apparent during the downstream trajectory analysis, where different cell types were disconnected, suggesting an over corrected batch effect (S1D, S2D). This was not the case for the human dataset, for which single-cell annotation using variational Inference (scANVI) performed the best (S2A). We therefore chose, to continue with scVI for integration and scANVI (Xu, 2021) for all cell type classification.

### Validation

To validate models we performed a series of downstream analyses from the learned scVI latent space (see Fig. 1C bottleneck layer annotated as Z). In both datasets, we started by computing the nearest neighbors graph (k-NNG; k=15), followed by Force Directed Graph (FA) and Uniform Manifold Approximation and Projection (UMAP) (McInnes et al., 2020) dimension reduction methods (S1F, S2F). Finally, we identified populations of cells using unsupervised Leiden clustering (Traag et al., 2019) and inferred differentiation trajectories using Partition-based graph abstraction (PAGA) (Wolf et al., 2019) (S1G, S2G).

In case of the mouse model, branching trajectories covering the first lineage decisions (TE, EPI and PrE) coincide with our understanding of *in vivo* development. We inspected the 15 identified clusters (resolution=0.8) and found, that the most dominant and heterogeneous populations was the ICM (S1C) which spans approximately five clusters (S1E). As the ICM is a very transient cell type and a progenitor of both the EPI and PrE, this behaviour is expected. Additionally, clustering was unable to distinguish between cleavage states with the 8C and 16C stages both contained in cluster 1, indicating that these stages are transcriptionally similar (S1E). PAGA trajectory inference correctly connected all developmental stages, confirming the underlying connected graph was consistent with development (S1G). To inspect gene activity, we inferred pseudotime, a proxy of developmental time using diffusion pseudotime (dpt) (Haghverdi et al., 2016) and scFates (Faure et al., 2023) package. Both tools were provided with the initial state set to zygote cells. While scFates and not dpt predicted a grade lineage transition reflecting the underling biology, both tools suffered from the same problem - they predicted the TE as a lagging lineage compared to EPI and PrE (S1H). However, we view this as a limitation of pseudotime being proxy and given that hierarchical clustering from scFates displayed the correct lineage segregation (S1I).

Generating a human model on the other hand, proved to be more complex as the scarcity of embryos meant a disproportionate quantity of specific cell types and the majority of the sampled cell types was originally annotated as TE cells (1,270 cells, S2B) as this cell type is the largest component of the human preimplantation embryo. As zygotic transcription in human does not start until the 8C, these cells collapsed together in FA graph, separated from the rest of the dataset (S2F) (Yuan et al., 2023). However, segregation of TE, EPI and PrE populations was confirmed using PAGA (S2G). Unsupervised clustering identified 18 clusters and was not able to distinguish individual cell types due to TE oversaturation (S2E). We used the same tools to predict pseudotime as with the mouse datasets. When pseudotime inference was applied with the ‘Prelineage’ cells as the initial state, dpt failed to assign any temporal sequence. scFates managed to infer a relatively correct pseudotime, but it failed to correctly assign terminal values to either EPI or PrE (S2H). Hierarchical segregation also failed in this instance as it identified TE 5.0 as a precursor to ICM and TE, suggest that more refinement is required. We hypothesize that refinement of the underlying scVI representation as well as identification of the ‘Unknown’ cells might align the model better with our understanding of human development.

### Classification

One of the most difficult and arduous tasks in scRNA-seq analysis is cell type classification; it requires an extensive knowledge of the studied system and is usually an intuitive process that is difficult to automate. Before training a computational classifier, we compiled and visualized a list of canonical markers from *in vivo* studies for each developmental stage (Fig. 1D) to ensure our starting labels were broadly correct. To overcome, accelerate and automate this task, we trained a series of machine learning classifiers based on either Gradient Boosting Decision Trees (GBDTs) and neural networks exploiting the tool we used in data integration. scANVI is a semi-supervised model which extends scVI by incorporating cell labels. In the background it refines the underlying scVI latent space while also learning the features that enables it to predict a cell type. By default, scANVI outputs the cell type classification with the highest score. For each cell, we use entropy as a measure of the uncertainty of this classification by subtracting this score from 1.0. GBDTs require a preexisting count matrix for training the cell type classifier. To generate this count matrix we used denoised RNA expression of HVGs generated from the scVI/scANVI and scGEN decoder for both species. Datasets were split 80:20 for training and testing of each cell type classification. We implemented this classifier using the XGBoost library. To mitigate overfitting, we allowed for early stoppage of the training if the logloss metric failed to improve in the previous 10 iterations. Lastly, we performed 10-fold cross validation to confirm the robustness of the classification model.

#### Mus Musculus

In mouse, the XGBoost classifiers performed the best in terms of the balanced accuracy metric with XGBoost[scVI] (0.96), XGBoost[scANVI] (0.91) and XGBoost[scGEN] (0.91) compared to scANVI (0.64) (Fig. 2A,S3A,B, Suppl. Table 2). The close to perfect performance of the XGBoost models suggested that they were overfit (Fig. 2A, S3A). scANVI performed relatively poorly (Fig. 2A, middle panel), but as scANVI is a neural net that looks at the proximity of all cells in latent space and our datasets are not large, the disproportionate representation of cell types could bias classification. In the case of the mouse, random samples would be more likely to select from the ICM and PrE than other cell types. To see if balancing the number of cells in the training set would improve classification, we trained a new scANVI (ns=15) model by setting 15 cells per cell type for each training epoch. This adjustment yielded a 23% increase in the balanced accuracy (0.87), strengthening the predictive power for E3.25-ICM/TE, E3.75-ICM and E4.5-TE which were previously misclassified. As in the clustering analysis, the most difficult cell type to predict was E3.5-ICM (N=457) with only a 46% prediction score. The remaining 54% of E3.5-ICM were predicted as TE, EPI, PrE and ICM of the E3.25 and E3.5 embryo (Fig. 2B, S3C). We believe the variability in ICM prediction is a result of the heterogeneous nature of ICM cells as they transition through the specification of EPI and PrE, prior to the segregation of these two lineages in time and space (Plusa et al., 2008; Ohnishi et al., 2013; Saiz et al., 2016).

**Figure 2.**
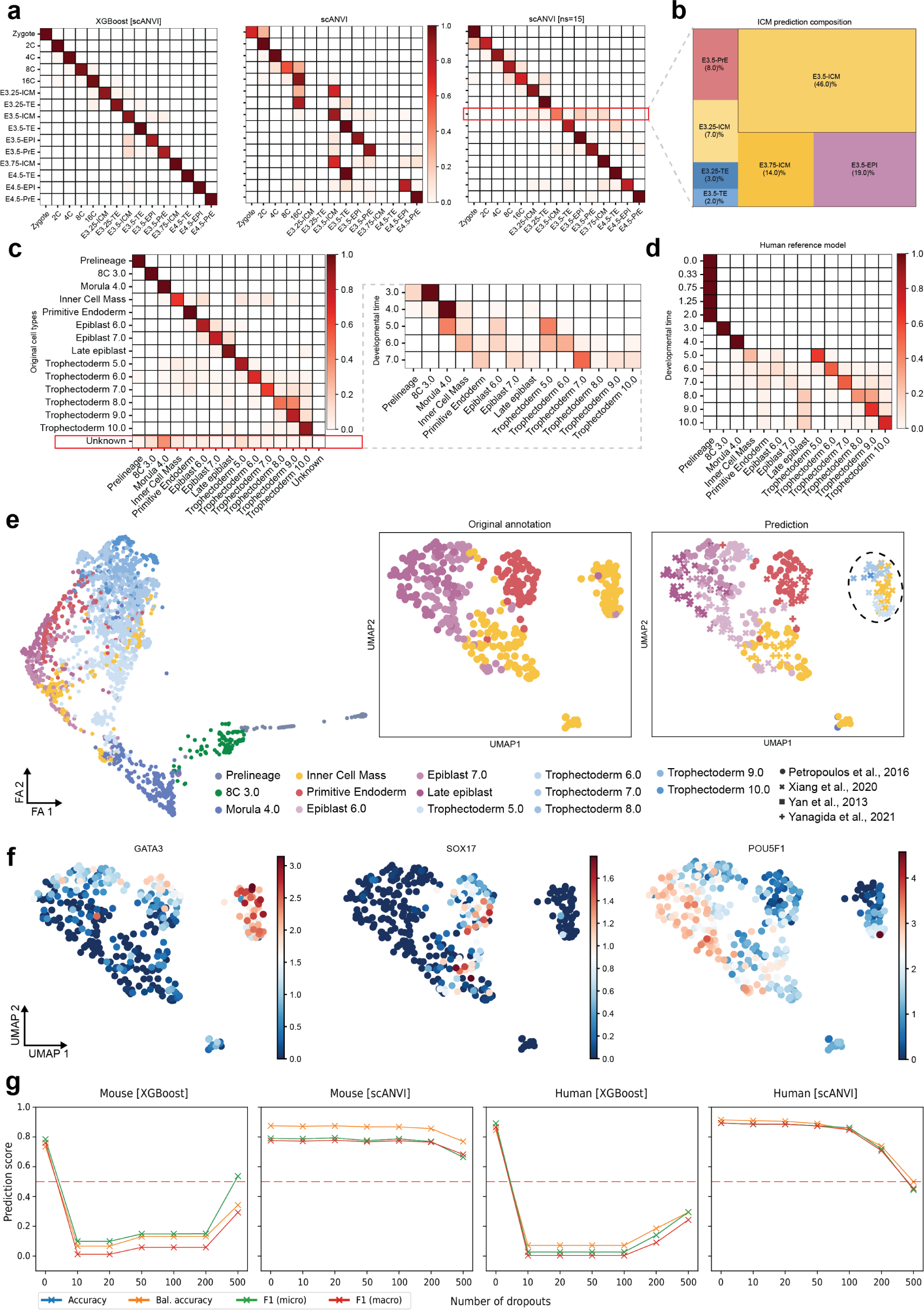
Cell type classification. **a**) Accuracy of predictions from three different mouse classifiers XGBoost (left), scANVI (middle) and scANVI with cell type sub-sampling (scANVI [ns=15], right). Scale represents prediction score for each individual cell type. **b**) Closer inspection of how annotated E3.5-ICM were predicted by scANVI[ns=15]. **c**) Accuracy of sub-sampled scANVI classifier (scANVI [ns=15], left) for the human reference, including reannotation of previously unannotated cells (right). **d**) Classifier annotations for cells sampled at known developmental times **e**) FA graph (left) and UMAP dimensional reduction (middle and right) displaying cells originally annotated as ICM but predicted to be trophectoderm. **f**) Expression of *GATA3*, *SOX17* and *POU5F1* in the ICM and ICM derivative subset. **g**) Impact of removing the top dispersion HVGs on classification performance of XGBoost and subset scANVI classifiers.

#### Homo Sapiens

Based on our observation with scANVI in mouse, we trained scANVI (ns=15) on a human dataset. As a result of this training exercise scANVI was now able to predict the identity of the ‘Unkown’ cells in the human dataset (Fig. 1E). The majority of unannotated cells at E3.0 and E4.0 were predicted by the scANVI to be cells of the prelineage embryo and morula, respectively (Fig. 2C left panel). To provide a better reference to the accuracy of these predictions they are represented alongside the time of collection in the right panel. Here the prediction for the stage of TE differentiation fits with the time point on which these cells were collected and thus suggests they were assigned to TE at the appropriate developmental age (Fig. 2C).

A second round of prediction with scANVI was performed on the total dataset and used to determine if any of previously classified cells might be misclassified (Fig. 2D). In third iteration, scANVI renannotatd a cluster of ICM cells towards early stages of TE (Fig. 2E). Fig. 2E (right panel) shows a subset of the complete dataset containing only cells from the ICM, EPI and PrE. Of the reannotated cells, a significant proportion originated from the Xiang et al. (2019) (Fig 2E,F), where other groups have observed inconsistencies in annotations (Zhao et al., 2021). Previous attempts at integrating preimplantation datasets have either omitted or manually reannotated this dataset (Meistermann et al., 2021; Zhao et al., 2021; Radley et al., 2023). As these cells do not cluster with the rest of ICM (Fig. 2E), express genes typically associated with the ICM, EPI (SOX2, NANOG, KLF4, KLF17 and POU5F1) (S3D) or those associated with PrE (SOX17, GATA4) (S3D), we have updated their annotation in our model based on the results obtained with scANVI.

#### Determining the XGBoost model is overfit

Based on the overfit of the classifications determined by the XGBoost-based classifier, we hypothesised the scANVI classifier would be more robust when applied to sparse and noisy datasets. To test this, we benchmarked the accuracy of both classifiers when they were provided with normalised gene expression after removing the top HVGs used to build them. Dropping as few as the top 10 HVGs reduced the performance of the XGBoost based classifier to close to 10% for all measured accuracies in both species (Fig. 2G, Suppl. Table 3). In both species, the balanced scANVI classifier was more robust to removal of the top HVGs, only significantly losing accuracy after removal of the top 200 HVGs. This is consistent with the functionality of XGBoost that predicts identity based on the cumulative score from each of the classifiers, making a sequence of binary judgment with the narrowest possible feature set. For scRNA-seq gene expression, where there is often stochastic variation of a single marker in a heterogeneous population (Elowitz et al., 2002), this approach would miss the nuances of cell type identification. The scANVI classifier takes advantage of integrating datasets into one latent space and then predicting cell types based on their proximity to known labeled cells, thus taking into an account a broad range of possible features. Based on these considerations, we chose to continue with scANVI [ns=15] for both species.

#### Explaining scANVI models

We next sought to understand what are the features that define a specific cell type; to uncover which genes were used to assign cell type identity and determine how these genes align with the known markers from the literature. In the case of XGBoost, one can extract deciding features (genes) used in overall classification, but not those for specific classes (cell types). scANVI is even more problematic as it suffers from a “black-box” issue given that its architecture is a neural network consisting of learned weights which are difficult to interpret. These shortcomings can be addressed with methods like SHAP (Strumbelj and Kononenko, 2010; Lundberg and Lee, 2017) or Local Interpretable Model-Agnostic Explanations (LIME) (Ribeiro et al., 2016) that test the importance of individual features (genes) for each prediction. Although SHAP has been applied to XGBoost, it has never been applied to neural networks like scANVI. We therefore devised a custom scANVI Explainer (scANVIExplainer), which estimates SHapley values to quantify the feature contributions in classifying predicted cell types.

To extract a robust feature set for each cell type, we bootstrapped the scANVIExplainer 10-times and discarded features which did not have positive weights through-out all 10 iterations (Fig. 3A). We then sought to assess the level to which these features made biological sense and performed differential expression analysis for identified features. Fig. 3B shows log fold gene expression changes for the top 3 ranked features as a heatmap. We provide a full list of features and their corresponding expected SHAP value in Suppl. Table 4 and S4.

**Figure 3.**
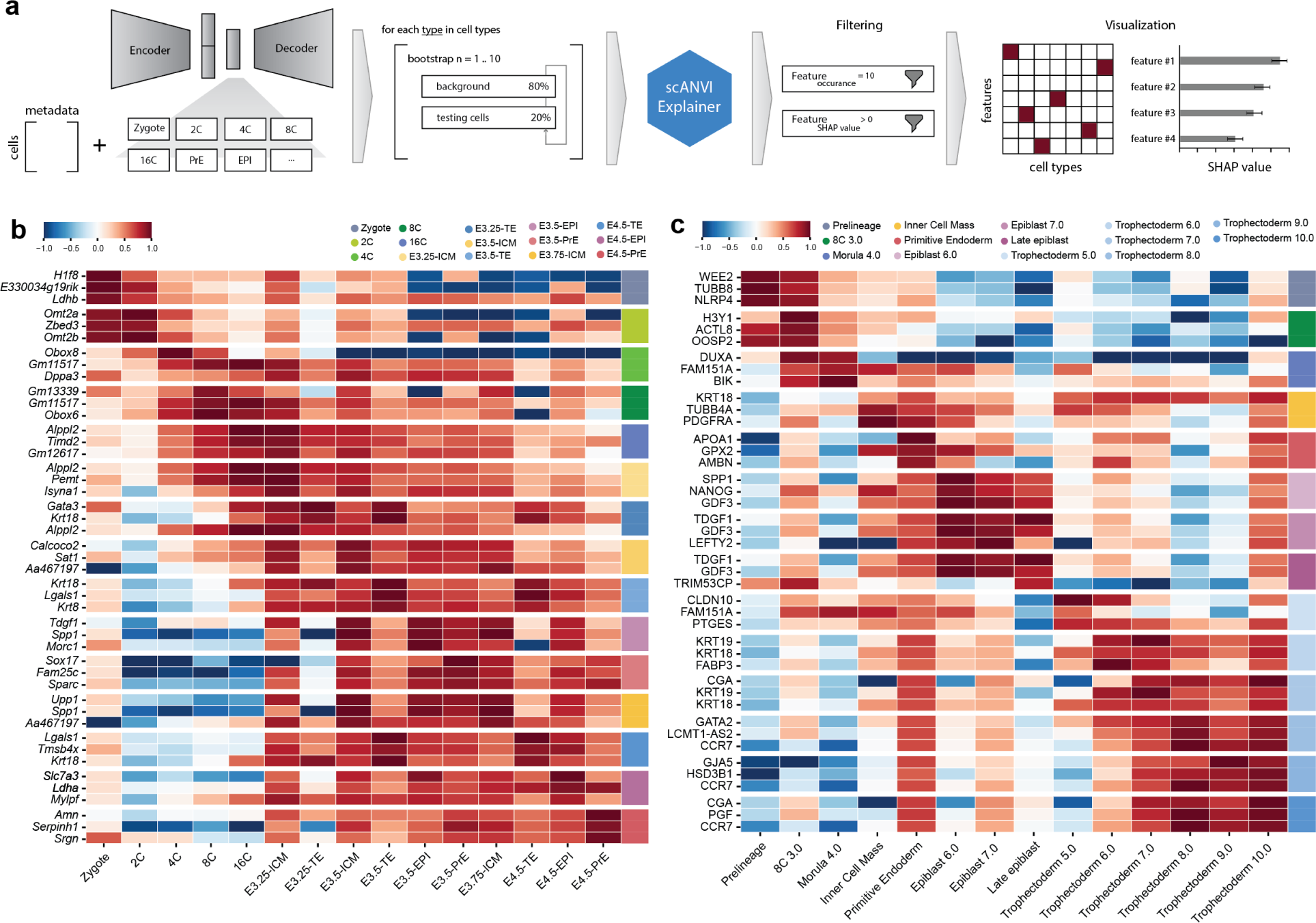
Extracting key predicting features with scANVIExplainer. **a**) Schematic overview of how scANVIExplainer works. **b-c**) Differential expression analysis (one vs all) of genes identified as top 3 predictors for each cell type in mouse (left) and human (right) classifiers. Heatmap displays Log_2_(fold change) of cell type vs all other cell types. Legend for vertical cell type identification is given at the top of each heatmap.

In both mouse and human, the classifier used a combination of canonical and non-canonical markers. In mouse, we found that some of the most prominent markers used for staining preimplantation lineages, such as *Cdx2* (TE), *Gata6* (PrE) and *Nanog* (ICM/EPI), were not reported in the top list. Instead we observed the model used genes associated with early development like *Omt2a*, *Obox8*, *Dppa3* alongside known markers like *Gata3* (TE), *Sox17* (PrE) and *SPP1* (ICM) at later stages of development. Similar analysis in the human model revealed the classification of transcriptomes to prelineage or 8C stages logically relied on the previously identified *NLRP* family gene *NLRP4* and the oocyte factor Oocyte secreted protein 2 (OOSP2). As with the mouse, the human scANVI model also utilised the expression of some traditional markers, such as the surface marker *PDGFRA* for PrE, the TGF-*β* ligands *NODAL* and *GDF3* for the EPI and several critical genes in placental development *KRT18*, *CGF* and *PGF* for TE (Suppl. Table 4, S4).

### Query integration: Annotating experimental datasets using the optimised scANVI models

One of the key advantages of the scANVI model is simple integration and classification of new datasets, showcasing that the models can accurately predict cell types generated *in vitro*. To test this, we explored two ESC derived cases of *in vitro* differentiation, one mouse and one human.

#### Mouse in vitro PrE differentiation

As a model system for mouse PrE differentiation we used an *in vitro* PrE differentiation protocol (Anderson et al., 2017). We recently published a single-cell MARS-seq dataset (Perera et al., 2022; Proks et al., 2023) for this protocol that exploited double fluorescent reporter cell line that contained fluorescent protein reporters for both an EPI or pluripotency marker Sox2 alongside an early marker for PrE (Hhex); Hhex-mCherry and Sox2-GFP. Reporter ESCs cells were passaged in defined naïve pluripotent media conditions, containing two small molecular inhibitors for GSK and MEK and the cytokine LIF (2i/LIF) in N2B27 media. For differentiation they were plated into an RPMI based differentiation media containing Activin, Chiron and LIF (RACL) for 7 days using the protocol we described previously (Anderson et al., 2017). Cells were collected at different stages of development, sorted and equal quantities of each population sequenced using MARS-seq (Jaitin et al., 2014). Based on imaging and cell sorting we had been able to define the PrE differentiation trajectory in the clusters of cells expressing HHEX ((Perera et al., 2022; Proks et al., 2023)), but now we were able to use scANVI to test this hypothesis and map this population onto *in vivo* developmental stages. Fig. 3B shows the dataset, flow cytometry and predicted cell types indicating that this trajectory passes through an E3.5 PrE state to a terminal state at E4.5 PrE, while the population that fails to differentiate retains an EPI-like identity. That the remaining SOX2+ cells were classified as E3.5/E4.5 EPI like cells, coinciding with our previous findings that these cells have gene expression signature that resembles pluripotency. Fig. 4A (Proks et al., 2023) also shows an entropy score (uncertainity) for the scANVI prediction, that suggests accurate predictions, particular at the end of the differentiation protocol (Fig. 4C left panel).

**Figure 4.**
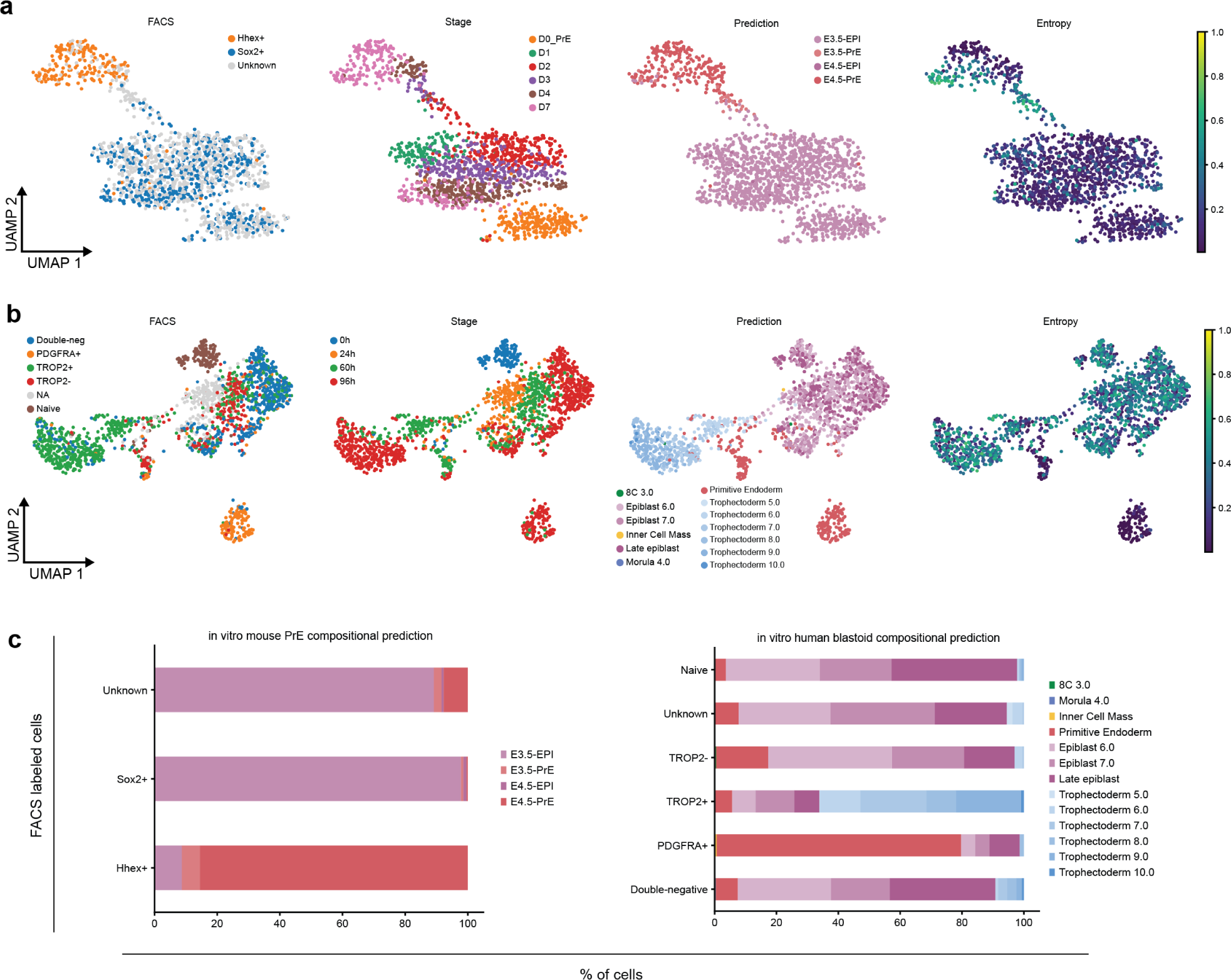
Classifying *in vitro* datasets. **a**) Prediction of cell types generated during mouse *in vitro* primitive endoderm differentiation in HHex/Sox2 double-reporter embryonic stem cells. **b**) Prediction of cell types generated in a human stem cell-based model of blastocyst development. **c**) Left panel: predicted cell type proportions within mouse primitive endoderm differentiation from embryonic stem cells compared to reporter expression; Right panel: predicted cell type proportions within *in vitro* human blastoids compared to cell surface marker expression.

#### Human naive stem cells and blastoids

To probe the capabilities of our human classifier to predict *in vitro* cell type identity we tested its ability to assess *in vitro* three dimensional models for early human development. A number of recent studies have demonstrated that naïve human ESCs (hESCs) are capable of self organizing into integrated stem cell-based models of human blastocysts called blastoids (Kagawa et al., 2021; Liu et al., 2021; Karvas et al., 2023; Yu et al., 2023). While scRNA-seq datasets have been generated from these structures in a range of conditions, we chose an embryonic stem cell derived model that had both RNA-seq and cell surface marker annotations from which we could test the capacity of our model to intuit cell identity (Kagawa et al., 2021). These cells were made from naïve hESCs cultured in PXGL (Bredenkamp et al., 2019b) (expand as above) and then transferred into microwells in conditions that promote blastoid formation, assessed by both flow cytometry for specific lineage markers (PDGFRA+ PrE; TROP2+ TE; Double neg EPI) and scRNA-seq. Fig. 4B shows the flow cytometry and time course for differentiation into blastoids in the first two panels, alongside the predicted scANVI identity and entropy. While specific predictions have lower certainty (higher entropy) than we achieved in mouse, scANVI reliably predicted PrE and TE maturation. As expected we find there is a large EPI-like population, that has a similar identity to the original naïve hESCs. While it has been suggested that naïve hESCs resemble early EPI and dedifferentiate to an ICM identity to generate TE in blastoids (Kagawa et al., 2021; Guo et al., 2021), scANVI does not predict any of the early time point blastoid cells to have an ICM identity. Taken together we demonstrated the capacity of our scANVI classifier to act in the absence of manual curration and an extensive knowledge of developmental biology to assign cell identities based solely on scRNA-seq transcriptomics (Fig. 4C).

## Discussion

In this paper we have shown that deep-learning tools can be applied to scRNA-seq from early embryos to generate dynamic models with predictive power that can be used to benchmark *in vitro* cell types. We do this from a readily assembled, compartmentalized set of tools wrapped in nf-core pipelines. We find that not only are these models capable of predicting cell types, but we can utilize our newly built scANVIExplainer to estimate SHapley values and produce a set of markers which can be used as a future basis to define cell types in an unbiased way. To facilitate the uptake of our model, we have developed a portal (https://brickman-preimplantation.streamlit.app) for inspecting and visualizing reference models. We also provide a Google Colab notebook containing code to easily retrain or query new datasets. We also invite others to augment the models with further reference datasets to strengthen the robustness of integration and cell type classification. We anticipate increased accuracy with expanded datasets as well as opportunities to benchmark current and new *in vitro* datasets.

Cell type classification has traditionally been based on morphology, functionality and position within an embryo or adult organism (Systems, 2017; Mulas et al., 2021; Fleck et al., 2023). In the wake of the molecular revolution in developmental biology and differentiation, identification became based on gene expression of select markers that were historically associated with specific cell types. Thus markers were discovered based on gene expression and functionality, and then came to define specific cell types. This has meant that lineages were defined based on specific markers such as *Nanog* or *Sox2* for EPI, *Cdx2* or *Eomes* for TE or *Gata4* and *Gata6* for PrE (Riveiro and Brickman, 2020; Biondic et al., 2023). As a result, most experimentally induced phenotypes have relied on this accumulated knowledge, rather than taking a systematic and unbiased approach to define cell type identity. Recent advances in our ability to capture whole genome expression in single-cells suggests RNA best describes a range of gene expression identities that comprise a cell state rather than discrete cell types (Huang, 2006; Enver et al., 2009). These cell states are defined using unsupervised clustering to produce populations of cells that are then annotated based on differential expression analysis and manual curation. Although there are known factors that will define a cell state, by taking an unbiased approach to defining the minimal set required to assign cell state identity, we find a mix of canonical and non-canonical factors. Thus, we did not observe canonical markers such as Cdx2/*Sox2*/*Gata6* as top ranked features enabling the discrimination of TE/EPI/PrE for which they are commonly used. Instead, our model used genes associated with early development like *Omt2a*, *Obox8*, *Dppa3* alongside *Gata3* (TE), *Sox17* (PrE) and *SPP1* (ICM) at later stages for mouse. Moreover, while our human classifier also avoided factors like *SOX2* and *GATA6*, it reiterated the importance of *PDGFRA* as a PrE marker, also highlighting the significance of TGF-*β* family genes in epiblast (*NODAL* and *GDF3*) and *NANOG*. Taken together, this suggests that although experimental developmental biology has identified some key markers, the accurate identification of cell types requires a more unbiased approach.

While our current models are limited to preimplantation development, as we used scANVI as the underlying architecture, we propose the model can still be used to benchmark *in vitro* cell types not contained in our dataset based on the entropy score (Xu, 2021). In both the human and the mouse, we observe naïve ESCs are EPI-like and are competent to form different extra-embryonic lineages. In human we did not observe any evidence that the cells regress to an ICM-like identity, but instead suggest they directly differentiate toward both extra-embryonic lineages. This is consistent with observations made over the years about the behavior of ESCs when reintroduced into development, where under certain conditions they can be observed colonizing all lineages (Beddington and Robertson, 1989; Macfarlan et al., 2012; Morgani et al., 2013; Gonzalez et al., 2016; Riveiro and Brickman, 2020; Redó-Riveiro et al., 2024).

The last year has seen the development of an abundance of *in vitro* stem cell-based model systems and revised *ex vivo* culture conditions for human embryos (Oldak et al., 2023; Weatherbee et al., 2023). The online resource accompanying this paper, while useful, highlights the need for new approaches to both classification and phenotype analysis. We believe our models are an excellent complement to already established large organ atlases (Regev et al., 2017; Swamy et al., 2021; Consortium* et al., 2022; Eraslan et al., 2022; Domínguez Conde et al., 2022; Suo et al., 2022; Sikkema et al., 2023) and attempts to model gastrulation (Nowotschin et al., 2019; Pijuan-Sala et al., 2019; Mittnenzweig et al., 2021). Given the scarcity and difficulty in obtaining human material, models like these represent essential resources for computational interrogation of genetic and biochemical perturbation of early development and *in vitro* models.

### Limitations of the study

One could argue that scANVI models are still limited by the usage of highly variable genes which is analogous to feature extraction. This is true, however both (Luecken et al., 2022; Xu, 2021) demonstrated that similar integrations using the full genome did not significantly improve overall performance.

One of the major the limitations of this study is the number of cells and imbalanced cell types. This is complicated by the non-trivial task of embryo dissection and the scarcity of human material. Despite this, our downstream analyses provides a snapshot of development that is consistent with existing knowledge of *in vivo* development. Despite the abundance of TE in our human dataset, our approach to balancing the classifier produced a reasonably high fidelity model. However, in the absence of experimental validation we cannot exclude the possibility that our existing classifiers are optimal identifiers of specific cell types. With time and augmentation of our model with new datasets, we believe that our classifier will continually improve, but the need for experimental validation is essential.

## Methods

### Downloading and preprocessing of single-cell RNA-seq data

We compiled a list of publication from numerous databases and publications (Svensson et al., 2020). Raw sequencing (FASTQ) files were downloaded from NCBI GEO and ENA repositories using the nf-core/fetchngs (v1.10.0) pipeline (Patel et al., 2023). For gene expression quantification, we forked nf-core/scrnaseq (Peltzer et al., 2023) to brickmanlab/scrnaseq (branch: feature/scrnaseq) and adapted it to support SMART-seq 1/2 experiments. Transcript abundance was estimated using STARsolo using indexes built from the GRCm38 reference genome and Ensembl 102 gene annotations or the GRCh38 reference genome and Ensembl 110 gene annotations for mouse and human datasets, respectively.

### Data harmonization and normalization

The metadata for each cell in the dataset contains information about the experiment/publication, sequencing technology, batches and original cell type annotation. We harmonized the original annotations as follows: (1) In the case of the mouse data, if a cell type was defined as 2C early/mid/late, these cells were reannotated to 2C cells. (2) In the case of the human data, where some cell types were transitionary, these were reannotated as “Unknown”. These reannotations were saved as “ct”, for cell type, in the metadata.

Gene transcript quantification was further processed using the scanpy (Wolf et al., 2018) library. To ensure compatibility between UMI based and full-length technologies, full length (SMART-seq) sequencing was normalised to mean gene length obtained from Ensembl gene annotations (GTF) using gtftools (Li et al., 2022) and rounded to the nearest integer. For mouse datasets, genes were filtered to exclude ribosomal, cell cycle, mitochondrial genes and those which were expressed in fewer than 10 cells. Cells were filtered to exclude those expressing greater than 20,000 genes and 26,000,000 counts. Raw counts were then depth normalised to median counts in the filtered dataset and Log1p transformed. For human datasets, genes were filtered to exclude ribosomal, mitochondrial genes and those which were expressed in fewer than 10 cells. Raw counts were depth normalised to 10,000 total counts and Log1p transformed.

### Integration and classification training

The top 3,000 highly variable genes (HVGs) were identified using the `sc.pp.highly_variable_genes` function within scanpy, with the following arguments: `flavour=“cell_ranger”` and `batch_key=“experiment”`. The scVI models were then built using 2 hidden layers and a negative binomial gene likelihood and trained for a maximum of 400 epochs, with early stoppage. The scANVI model was built on top of the scVI integration with provided cell type labels and trained for 15 epochs. scGEN was trained using normalized counts, with `batch_key=“batch”` and `labels_key=“ct”`, for a maximum of 100 epochs, with a batch size of 32 and early stoppage enabled. To compare PCA, scVI, scANVI and scGen integration methods, we used the GPU-accelerated scib-metrics python package (v0.4.1) (YosefLab/scib-metrics, 2023) to compute the evaluation metrics defined in (Luecken et al., 2022).

To fine tune the arguments used to build the reference models, we took advantage of an experimental scvi.autotune package which allowed for rapid model generation using combinations of arguments. In both datasets we adjusted the search space with the following parameters: gene_likelihood (nb, zinb), gene dispersion (gene, gene-batch), number of hidden layers (128, 144, 256), number of layers (2-5) and learning rate range (between 1e-4 and 0.6). For each hyper parameter optimisation, we measured the validation loss with a maximum of 100 training epochs and generated, in total, 50 models (Suppl. Table 1).

The gradient boosted decision tree classifiers were built using the XGBoost library, trained on denoised expression matrices obtained using normalized expression values from the scVI, scANVI and scGen models: `get_normalized_expression` function (return_mean=True, return_numpy=True). Predictions were benchmarked by calculating accuracy, balanced accuracy, f1 (micro and macro) scores using the scikit-learn python package (Pedregosa et al., 2011), where the test set was the whole reference model.

### Dimensional reduction visualisation and trajectory inference

Nearest neighbour graphs (k-NNG, k=15) were calculated from the scVI normalized learned latent space (`get_latent_representation` function) using the scanpy function `sc.pp.neighbors`. To identify cell clusters, we used unsupervised Leiden clustering (Traag et al., 2019) with resolution set to 0.8. Next, we generate Principal Component Analysis to inspect visually spread of cell type. Using the k-NN graph, we performed dimension reduction using UMAP (McInnes et al., 2020) and Force Directed Graph (FA) to visualize data in a 2D plot.

Trajectory inference was performed by calculating a diffusion map (sc.tl.diffmap) followed by PAGA analysis (Wolf et al., 2019) (sc.tl.paga) with default settings. Pseudotime was estimated using the dpt (Haghverdi et al., 2016) package, with “Zygote” specified as the initial state for both species. For scFates (Faure et al., 2023), we recomputed diffusion map and found the optimal sigma value (500 for both) using the `scf.tl.explore_sigma` function. Based on the scFates visualization `scf.pl.graph`, the initial states were set to “Zygote” and “4C” for mouse and human respectfully. Estimated pseudotime was scaled from 0 to 1 as is reported by the dpt package.

### scANVI explainer

To explain which features (genes) are used to determine a cell type we used SHAP (SHapley Additive exPlanations) (Lundberg and Lee, 2017; Lundberg et al., 2020). For XGBoost we use `shap.GPUTreeExplainer`. For scANVI we had to modify the original DeepExplainer because scANVI requires count matrix (X), batch and cell type annotation (labels) for initialization. We next modified code to call classifier for the learned latent space (Z). We provide the code for the scANVI explainer in `deep_scanvi.py`. For all classifiers, we split the reference dataset to 80:20 (background, test dataset) to obtain an expected value for each feature. This is executed 10 times and we keep only the features which occurred 10 times. Next, we subset for non-negative features only and calculate mean and standard deviation for each of them. Lastly, we rank the features for each predicted cell type.

### *In vitro* predictions

For the mouse *in vitro* dataset, we downloaded the raw count matrix from (Proks et al., 2023) and proceeded without reprocessing. The human blastoid scRNA-seq dataset was reanalysed from raw sequencing reads. The list of accession numbers was fed into nf-core/fetchngs which downloaded raw FASTQ files. These were then immediately preprocessed using the brickmanlab/scrnaseq pipeline. As the human dataset was sequenced using SMART-seq2, we normalized the raw count by mean gene length. Both datasets were integrated to their species appropriate scANVI reference model with the following settings: `max_epochs=100; plan_kwargs=dict(weight_decay=0.0) and check_val_every_n_epoch=10`. For each integrated cell, we obtained the predicted cell type, defined as the cell type with the highest probability. Uncertainty (entropy) for the prediction was calculated as 1 - the highest cell type probability: `1` `-lvae_q.predict(soft=True).max(axis=1)`.

## Data and code availability

Reference downloading and preprocessing pipelines can be found at https://github.com/nf-core/fetchngs (revision 1.10.0) and https://github.com/brickmanlab/scrnaseq (revision: `feature/smartseq`), respectively. Data analysis notebooks were uploaded to https://github.com/brickmanlab/proks-salehin-et-al. Portal was uploaded to https://github.com/brickmanlab/preimplantation-portal and deployed to https://brickman-preimplantation.streamlit.app. Dataset and trained models with parameters were uploaded to Hugging Face https://huggingface.co/brickmanlab.

## ACKNOWLEDGEMENTS

We thank Kathy Niakan, Sophie Petropoulos and Fredrik Lanner for valuable comments and feedback. We thank M. Linneberg-Agerholm for providing markers for identifying lineages in preimplantation embryos. Lastly, we thank Molly P. Lowndes, Marta Perera and M. Linneberg-Agerholm for proof reading the manuscript and members of the Brickman lab for constructive feedback on the analysis and the manuscript.

## AUTHOR CONTRIBUTIONS

M.P., N.S. and J.B wrote the manuscript. The computational work has been done by both M.P and N.S.

## Supplementary Information

**Figure S1.**
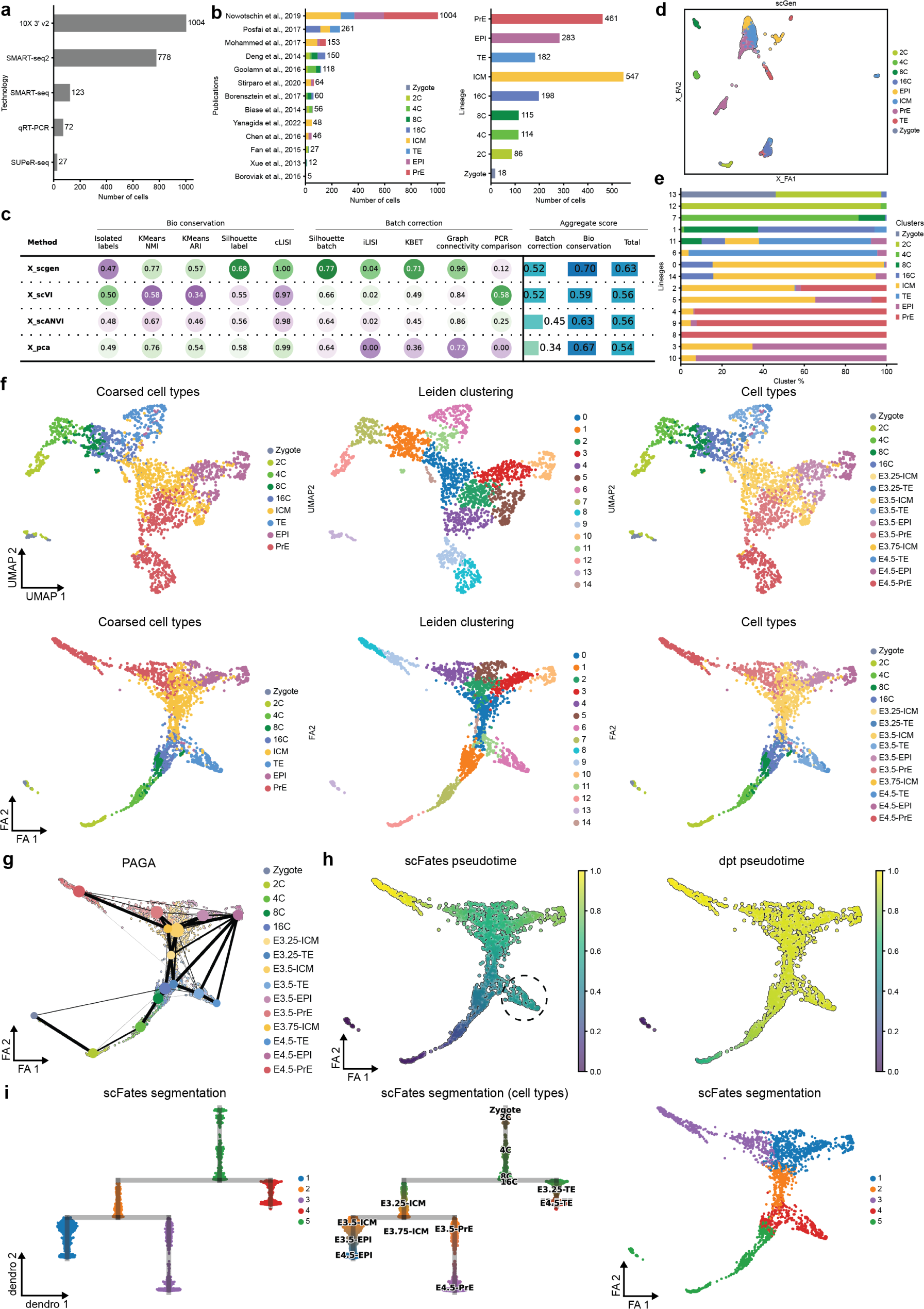
Mouse integration. **a**) Number of cell types per sequencing technology. **b**) Number of cell types per dataset (left) and lineage (right). **c**) Integration metrics for individual methods. **d**) Force directed graph of latent space inferred using scGEN. **e**) Cell type proportion per identified unsupersived clustering. **f**) Visualization of mouse reference dataset using UMAP (top row) and Force Directed Graph (bottom row) dimensional reduction techniques. **g**) Trajectory inference using PAGA. **h**) Pseudotime inference using scFates (left) and dpt (right) algorithms. **i**) Hierarchical clustering (left, middle) of segmentation (overlaid on Force directed graph representation, right) based on pseudotime from scFates.

**Figure S2.**
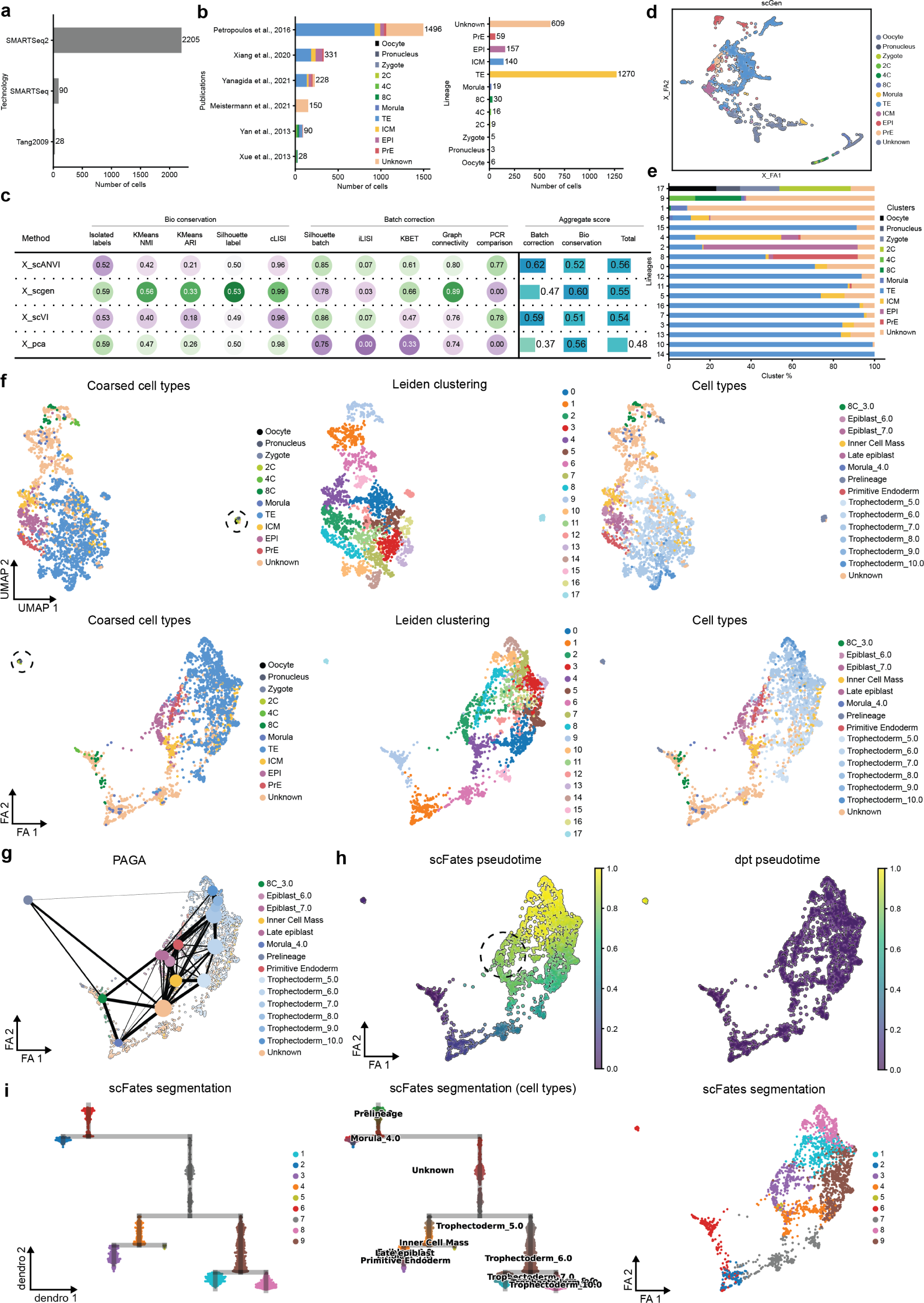
Human integration. **a**) Number of cell types per sequencing technology. **b**) Number of cell types per dataset (left) and lineage (right). **c**) Integration metrics for individual methods. **d**) Force directed graph of latent space inferred using scGEN. **e**) Cell type proportion per identified unsupersived clustering. **f**) Visualization of human reference dataset using UMAP (top row) and Force Directed Graph (bottom row) dimensional reduction techniques. **g**) Trajectory inference using PAGA. **h**) Pseudotime inference using scFates (left) and dpt (right) algorithms. **i**) Hierarchical clustering (left, middle) of segmentation (overlaid on Force directed graph representation, right) based on pseudotime from scFates.

**Figure S3.**
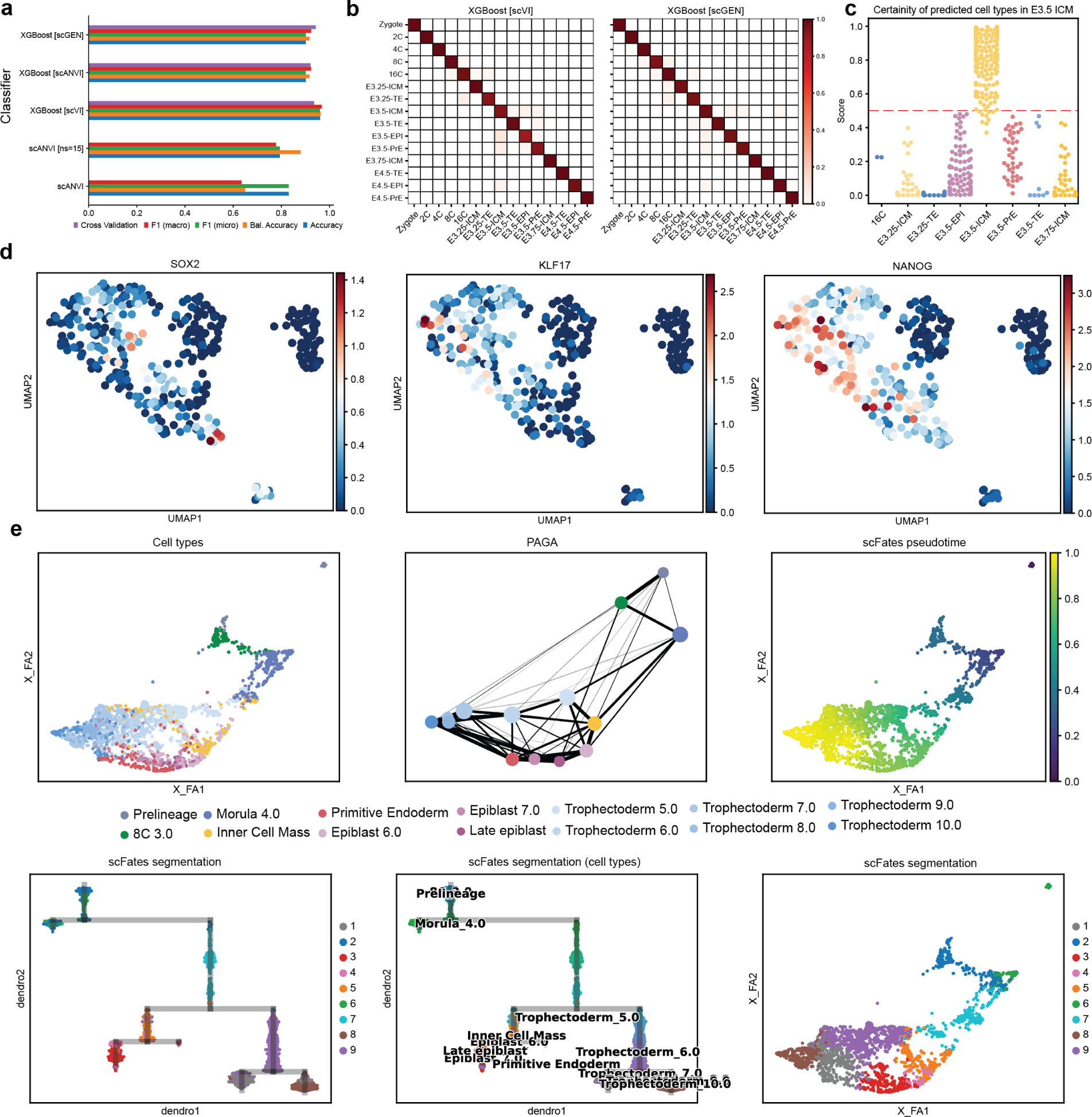
Classification metrics. **a**) Accuracy metrics for individual classification algorithms. **b**) Confusion matrix of XGBoost predictions based on denoised expression from either scVI (left) or scGEN (right) models, where y-axis are original annotations and x-axis are predictions. **c**) Maximum certainty of predictions of cells originally annotated as E3.5 ICM in the mouse dataset. **d**) Expression of *SOX2*, *KLF17* and *NANOG* in human ICM cells reannotated to trophectoderm using the scANVI classifier. **e**) Force directed graph visualisation of the human dataset after reannotation.

**Figure S4.**
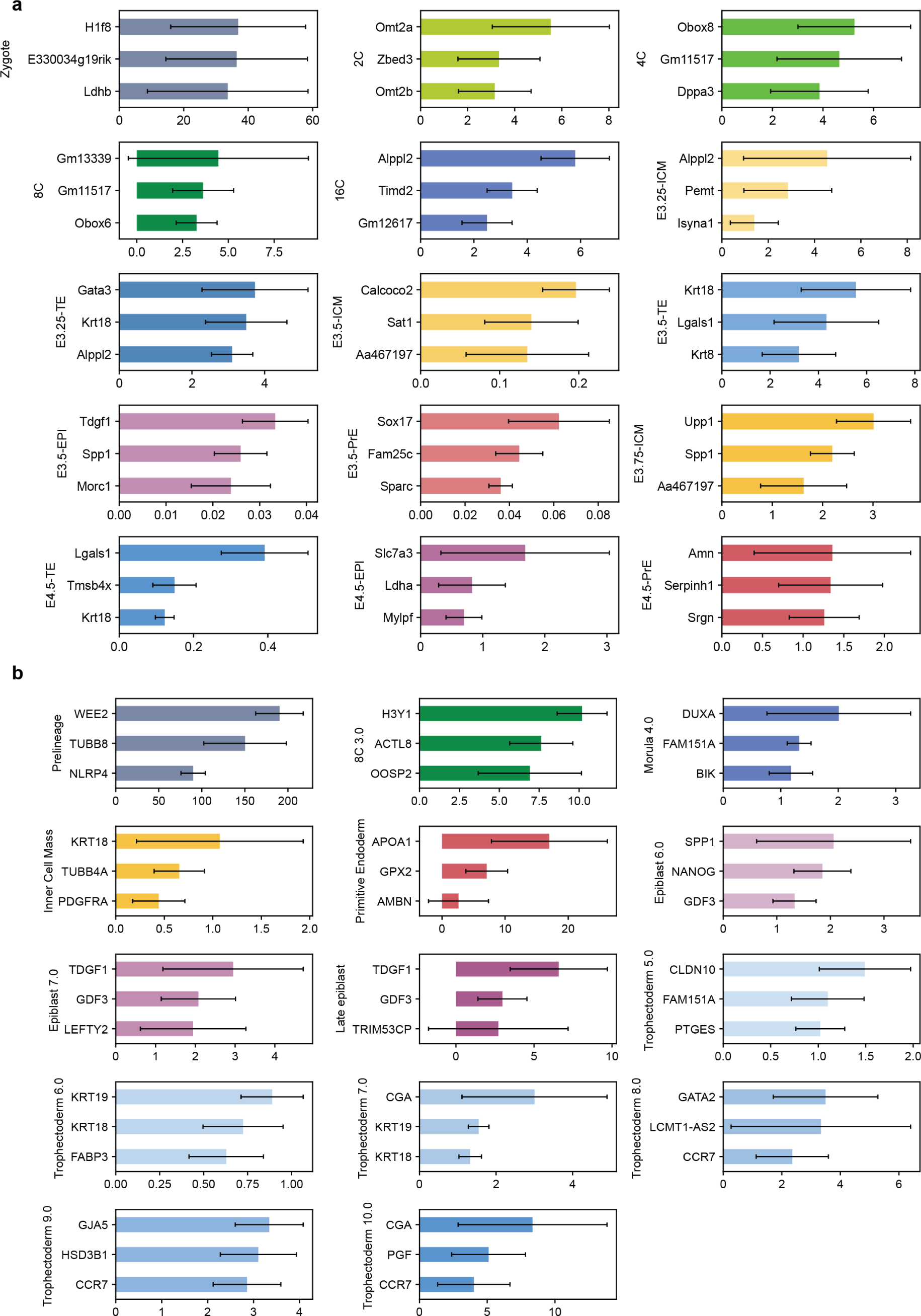
Feature importance for scANVI classifiers. **a**) Importance of genes in classification accuracy of individual cell types for the mouse scANVI classifier. **b**) Importance of genes in classification accuracy of individual cell types for the human scANVI classifier. Error bars represent mean ± standard deviation after ten randomised runs.

### Supplementary Tables

Table S1. Measured loss for different settings of scVI for mouse and human dataset.

Table S2. Accuracy metrics of predicting reference model for both mouse and human.

Table S3. Accuracy scores measuring resilience of scANVI and XGBoost predictions after removal of 0, 10, 20, 50, 100, 200 top ranked highly variable genes.

Table S4. All non-negative SHAP features extracted from scANVI after 10 bootstraps with confidence interval for both mouse and human reference model.

Table S5. Confusion matrix of mouse in vitro PrE differentiation and human in vitro blastoid differentiation cell type predictions.

